## Supplementary material for "Evaluating species interactions as a driver of phytophagous insect divergence": Figure S

SUPPLEMENTARY FIGURES AND TABLES

**Bruno A. S. de Medeiros**
Smithsonian Tropical Research Institute and
Museum of Comparative Zoology, Harvard University


**Brian D. Farrell**
Museum of Comparative Zoology, Harvard University


#### Contents

**1 Supplementary Figures 2**

**2 Supplementary Tables 10**

### 1 Supplementary Figures

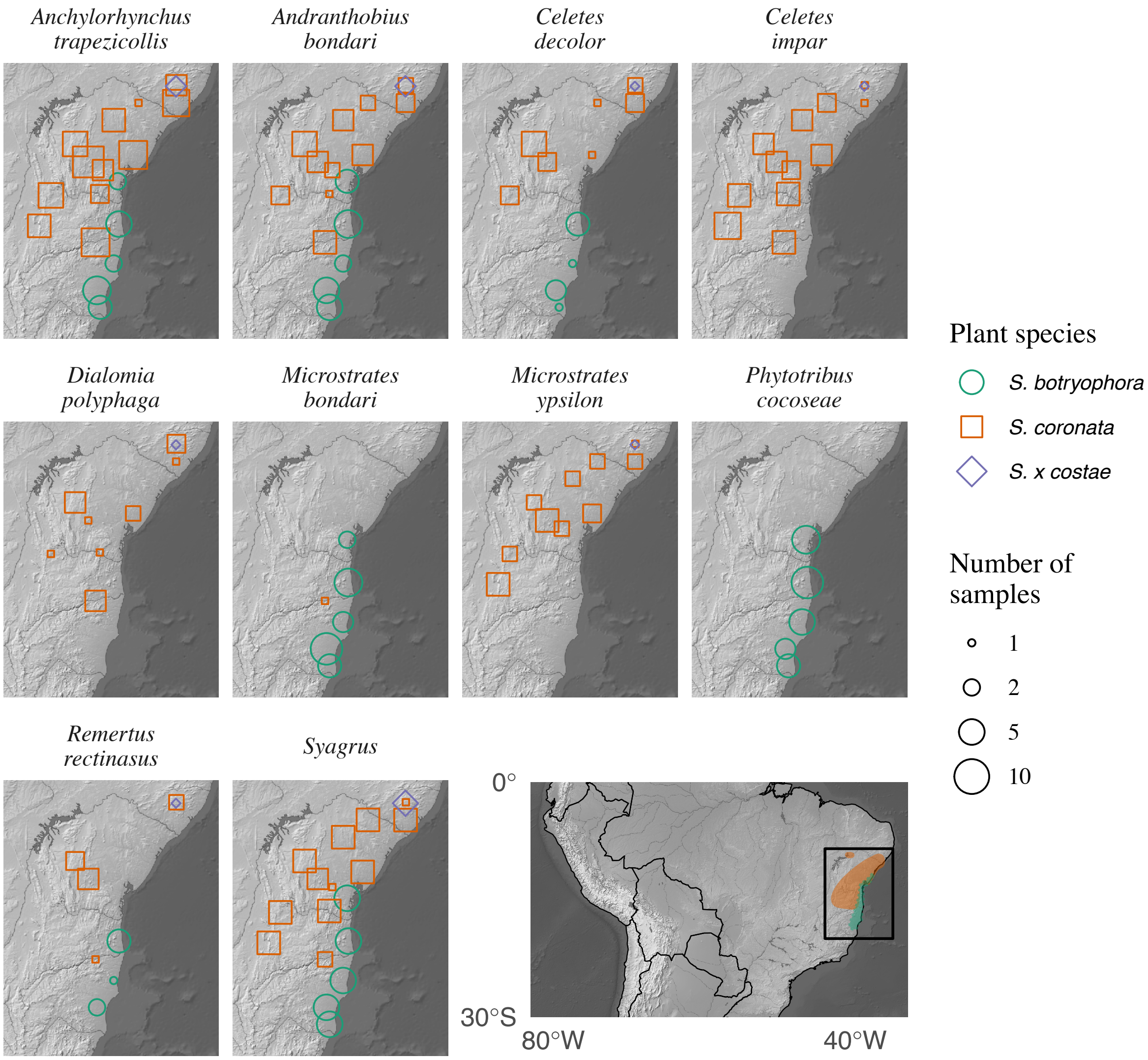


Figure S1: Sampling for species of palms and weevils. Large map shows the known distribution of palm species [2, 1]

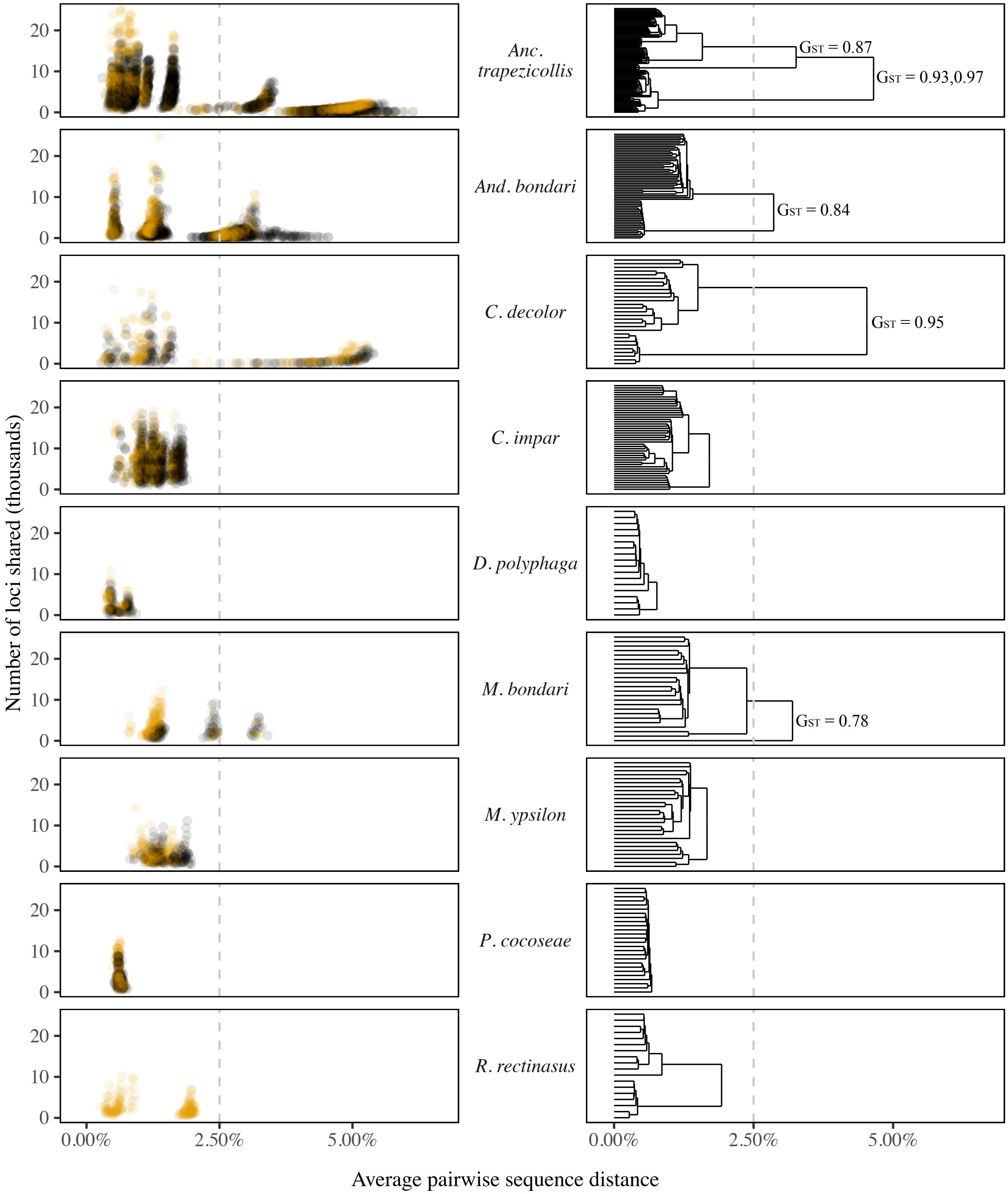


Figure S2: Left: pairwise sequence distance against pairwise number of loci shared. Yellow points show samples prepared in the same batch. Right: UPGMA clustering of samples based on pairwise sequence distance. Dashed line separates clusters at 2.5% divergence, with $G'_{ST}$ between these clusters shown.

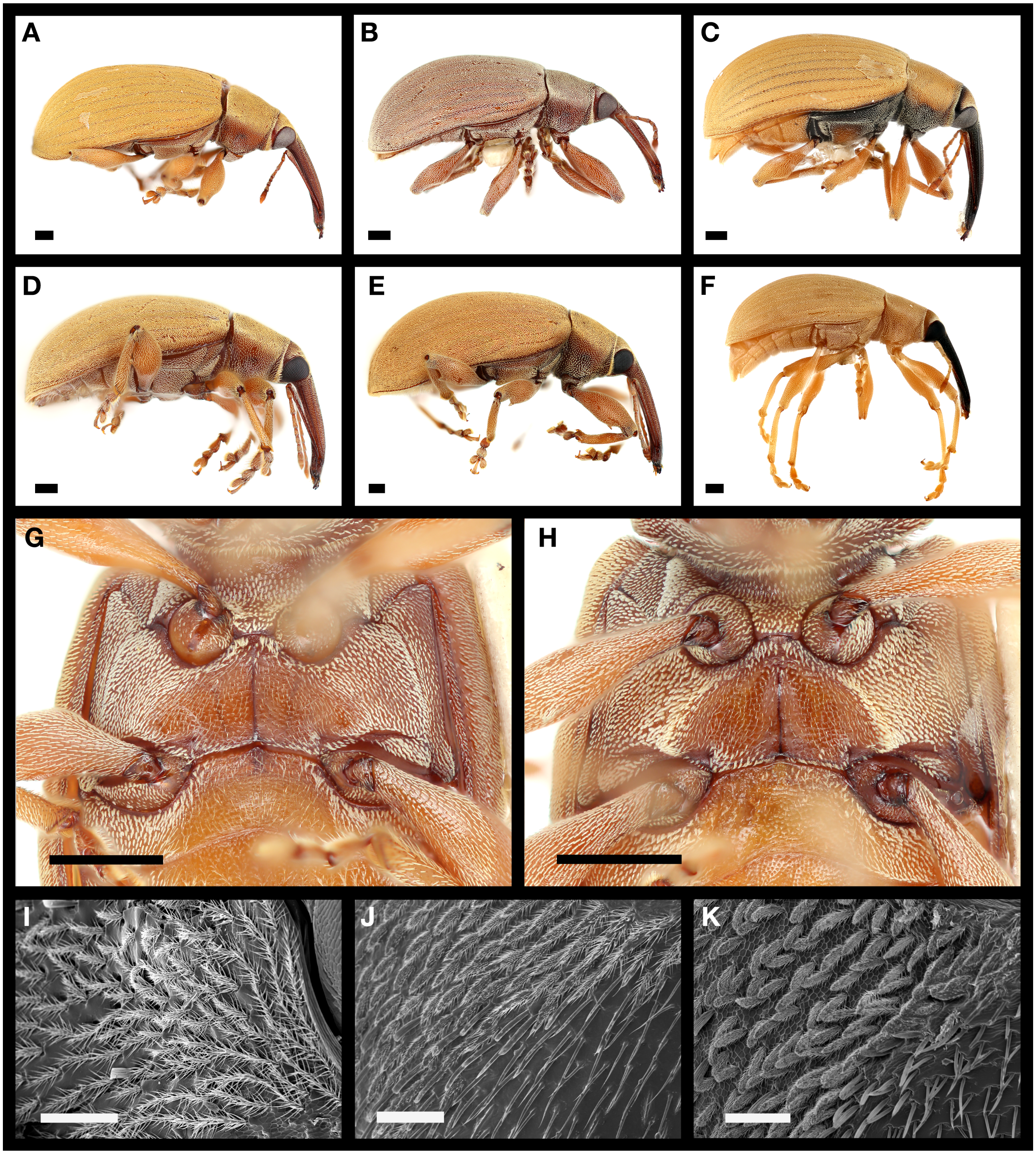


Figure S3: Morphology of *Anchylorhynchus trapezicollis* OTU 1 (**A–D,G,I–J**), OTU 2 (**E,H,K**) and OTU 3 (**F**). Black scales: 500 $\mu$m, white scales: 100 $\mu$m. See Figure 1 in main text for cluster labels. **A**: OTU 1, cluster C, **B**: OTU 1, cluster A (east), **C,G**: OTU 1, cluster A (west). **D**: OTU 1, cluster B. **E,H**: OTU 2, cluster B. Specimens **D–E,G–H** were collected in the same inflorescence. **I**: Ventral plumose setae in OTU 1, cluster B (in *Syagrus botryophora*) are very elongated and branched. **J**: Border of male metasternal concavity in OTU 1, cluster A (in *Syagrus coronata*), showing shorter setae in these populations. **K**: Border of metasternal concavity in OTU 2, cluster B, even shorter and less branched setae in this species.

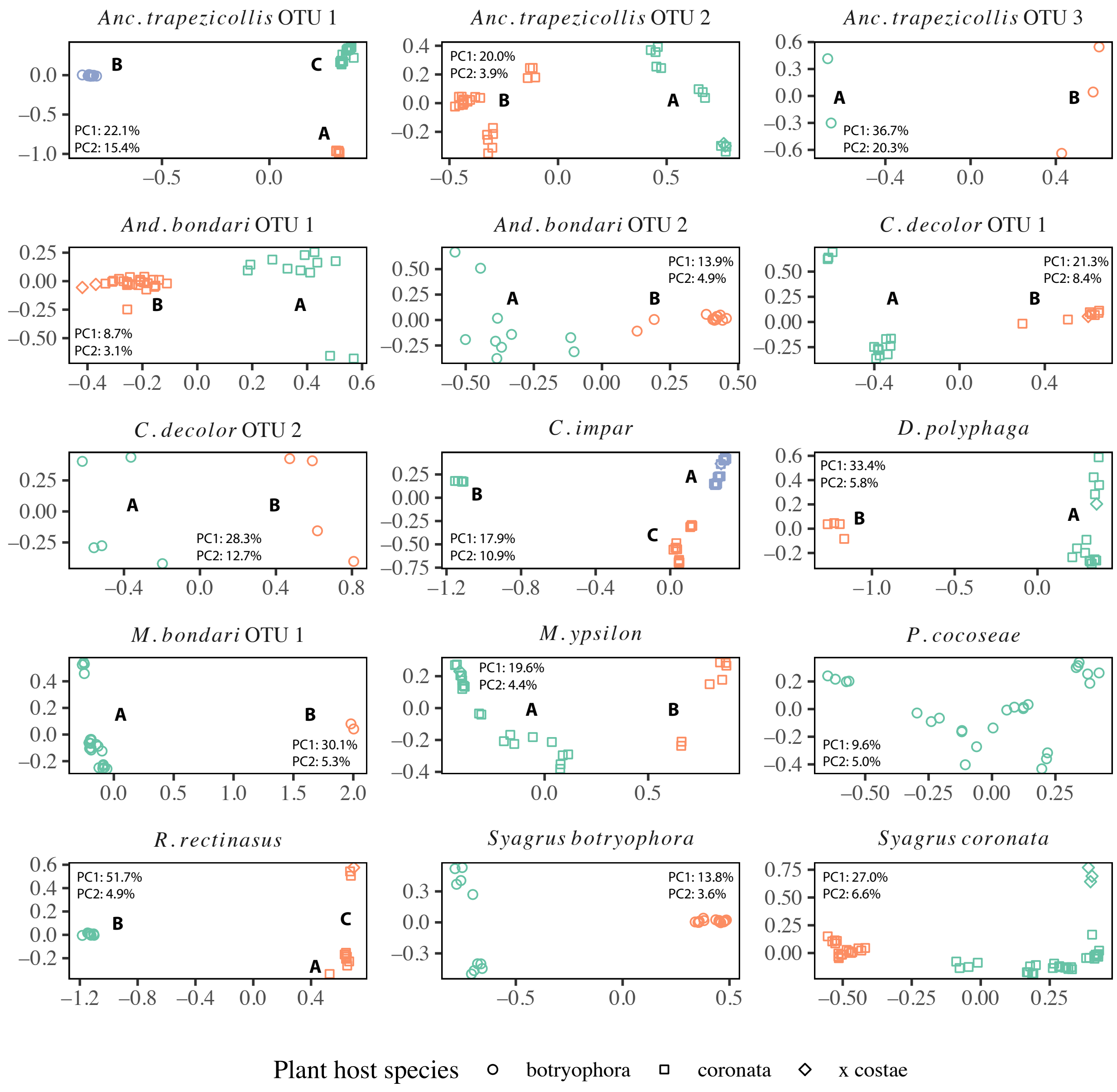


Figure S4: Principal component analyses of the genetic variation of weevil OTUs. Independently for each graph, different colors represent different clusters after k-means clustering. Percent variance associated with PC1 and PC2 are provided for each species. Cluster labels (A, B, C) correspond to populations used in coalescent analyses and also to Figure 1 in main text.

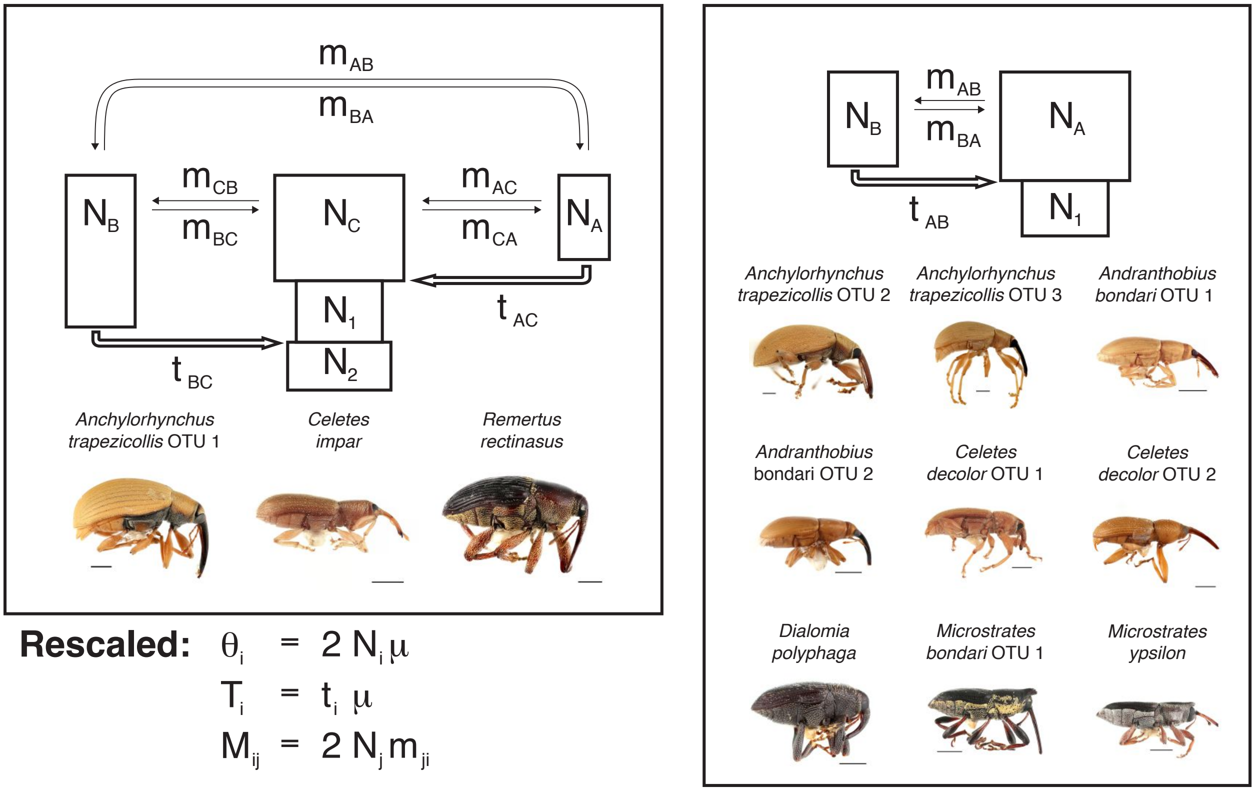


Figure S5: Full model of isolation-with-migration for OTUs with 3 or 2 populations. Boxes represent current (A, B, C) and ancestral (1, 2) populations. Thin arrows represent migration and thick arrows past population merge (backwards in time). Parameters inferred with fastsimcoal using a mutation rate ($\mu$) of 3e-9 were: **N** population sizes, **t** time to population merge (backwards in time) in generations, **m** migration rates (backwards in time). Inferred parameters were scaled to be independent of the unknown mutation rate using the equations in the figure.

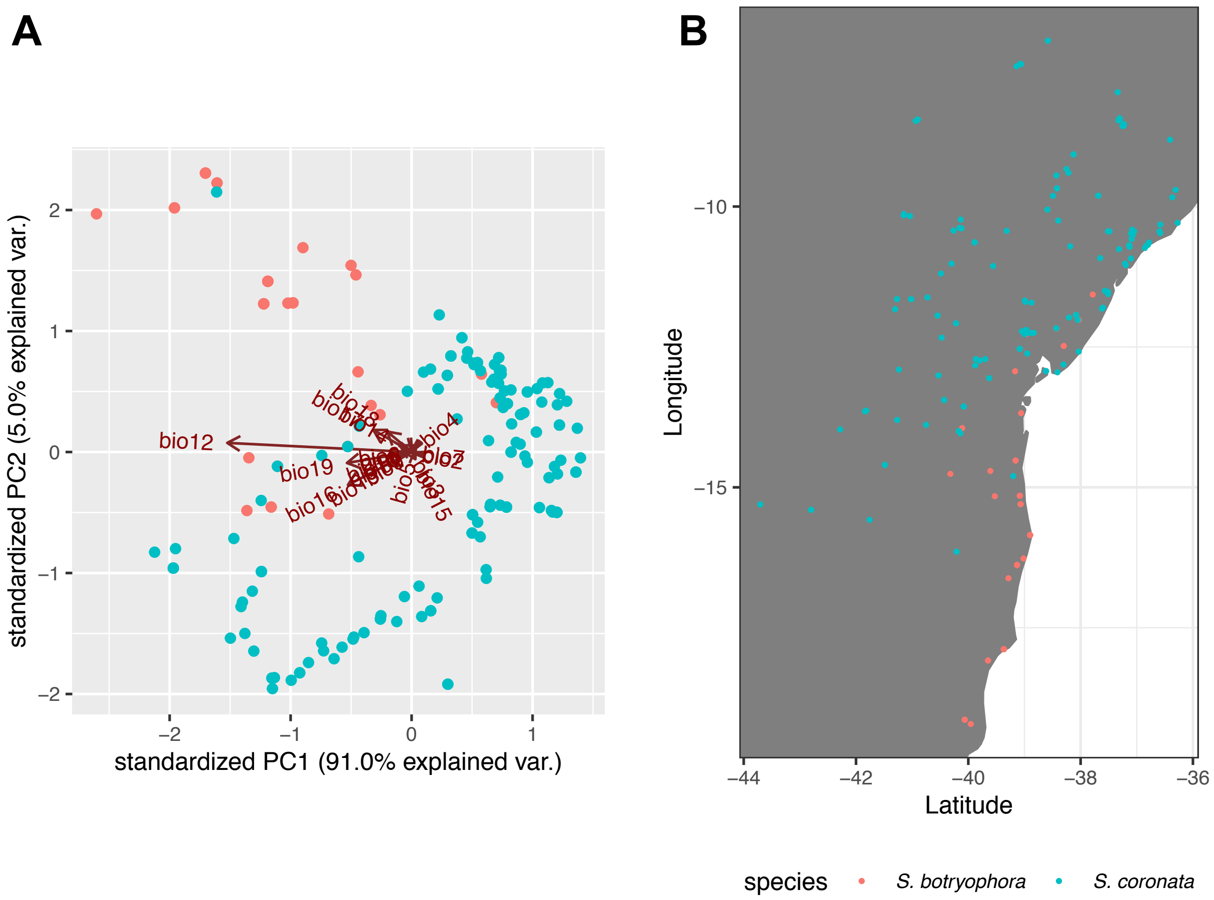


Figure S6: Annual precipitation explains most of the variation in climate across the distribution of both palms. **A** Principal component analysis of all bioclimatic variables extracted for occurrences of *Syagrus botryophora* and *Syagrus coronata*, showing the very high loading of bio12 (Annual Precipitation) on PC1, which explains over 90% of the variance . **B** Map of occurences used for this analysis.

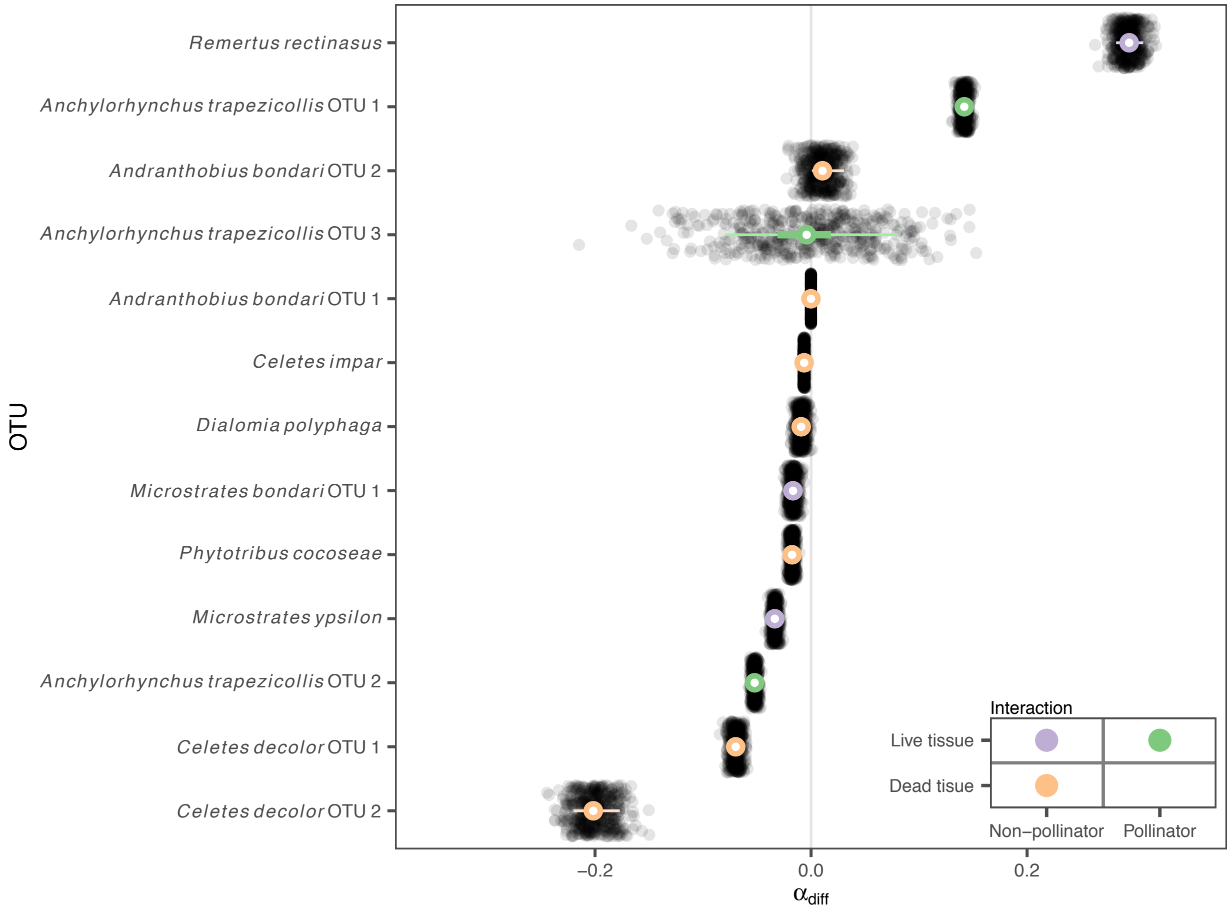


Figure S7: Posterior predictive simulations (points) show that the Bayesian hierarchical model has a good fit to the estimates of $\alpha_{diff}$ across species (colored ranges).

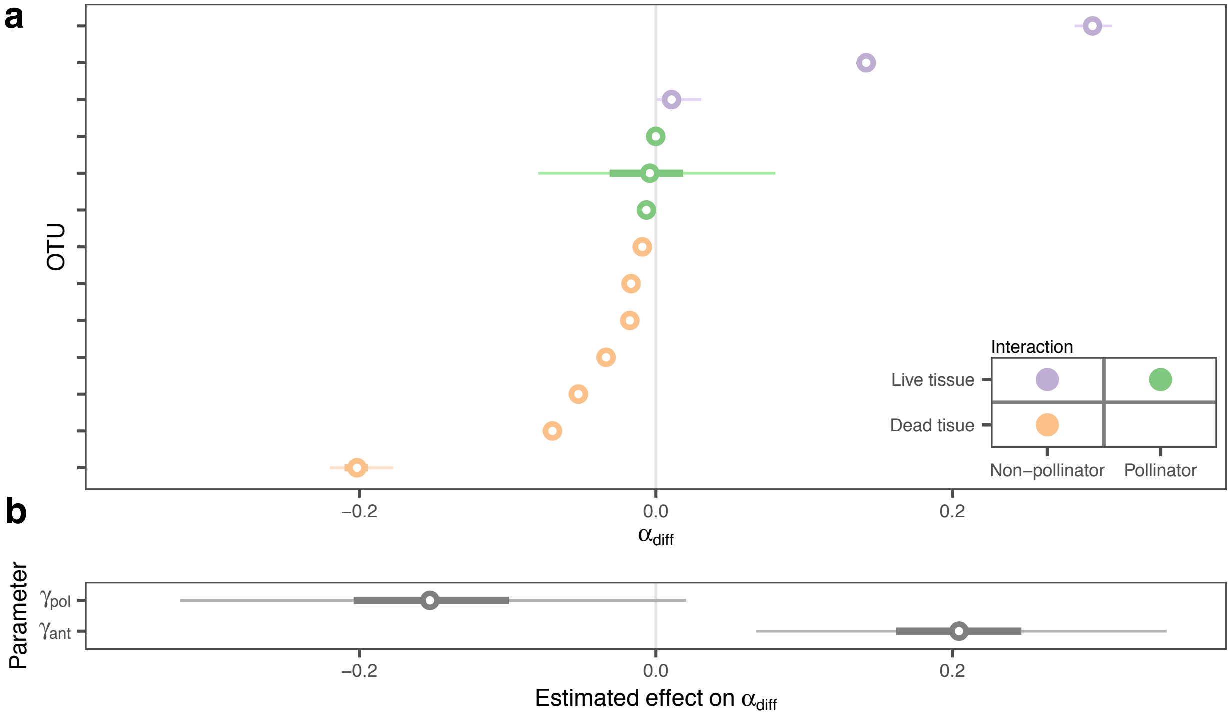


Figure S8: Relabelling of OTUs shows that the sample size and input data distribution in this study provide enough power to estimate $\gamma$ parameters. **a** Relabelled input data used as model input. **b** Estimated $\gamma$ parameters for this data.

#### 2 Supplementary Tables

Table S1: Models fitted in fastsimcoal, ordered by AIC scores and including maximum likelihood estimates. See Figure S5 for a visual scheme of model parameters. Preferred model for each OTU has the smallest AIC.

| OTU | Model | AIC | ΔAIC | $\theta_A$ | $\theta_B$ | $\theta_C$ | $\theta_1$ | $\theta_2$ | $T_1$ | $T_2$ | $M_{AB}$ | $M_{BA}$ | $M_{AC}$ | $M_{CA}$ | $M_{BC}$ | $M_{CB}$ |
|---|---|---|---|---|---|---|---|---|---|---|---|---|---|---|---|---|
| *Anchylorhynchus* | full | 481206 | 0 | 3.31E-03 | 3.69E-03 | 1.17E-02 | 2.15E-03 | 1.39E-03 | 3.61E-03 | 8.71E-03 | 0.07 | 0.05 | 0.19 | 0.49 | 0.15 | 0.18 |
| *trapezicollis* OTU 1 | no A,B | 483054 | 1848 | 3.32E-03 | 3.59E-03 | 1.19E-02 | 2.67E-03 | 4.00E-03 | 3.32E-03 | 6.19E-03 | | | 0.39 | 0.23 | 0.22 | 0.21 |
| | no A | 485764 | 4558 | 3.49E-03 | 3.51E-03 | 1.28E-02 | 5.06E-03 | 4.06E-03 | 1.84E-03 | 5.49E-03 | | | | | 0.23 | 0.17 |
| | no B,C | 486572 | 5366 | 3.46E-03 | 3.85E-03 | 1.25E-02 | 1.37E-03 | 5.56E-03 | 2.98E-03 | 3.66E-03 | 0.08 | 0.1 | 0.04 | 0.88 | | |
| | no B | 488324 | 7118 | 3.32E-03 | 3.97E-03 | 1.19E-02 | 1.23E-03 | 6.30E-03 | 2.66E-03 | 2.98E-03 | | | 0.17 | 0.36 | | |
| | no C | 488906 | 7700 | 3.32E-03 | 3.70E-03 | 1.60E-02 | 3.99E-03 | 5.80E-03 | 1.87E-03 | 3.37E-03 | 0.03 | 0.09 | | | | |
| | no mig | 489805 | 8599 | 3.27E-03 | 3.62E-03 | 1.40E-02 | 2.44E-03 | 6.35E-03 | 1.86E-03 | 2.62E-03 | | | | | | |
| *Anchylorhynchus* | full | 428578 | 0 | 3.32E-03 | 4.30E-03 | | 7.96E-04 | | 6.45E-03 | | 0.99 | 1.15 | | | | |
| *trapezicollis* OTU 2 | no mig | 434224 | 5645 | 3.27E-03 | 4.65E-03 | | 5.29E-03 | | 5.70E-04 | | | | | | | |
| *Anchylorhynchus* | full | 74284 | 0 | 1.42E-01 | 3.06E-03 | | 1.85E-02 | | 4.00E-03 | | 15252.22 | 0.27 | | | | |
| *trapezicollis* OTU 3 | no mig | 74512 | 228 | 3.54E-03 | 3.24E-03 | | 6.69E-03 | | 3.04E-05 | | | | | | | |
| *Andranthobius* | full | 637902 | 0 | 6.62E-03 | 1.79E-02 | | 3.51E-03 | | 3.70E-03 | | 9.6 | 21.46 | | | | |
| *bondari* OTU 1 | no mig | 644175 | 6273 | 2.03E-02 | 5.23E-02 | | 7.91E-03 | | 5.87E-04 | | | | | | | |
| *Andranthobius* | full | 350204 | 0 | 3.47E-03 | 8.55E-03 | | 1.11E-03 | | 5.35E-03 | | 4.21 | 9.78 | | | | |
| *bondari* OTU 2 | no mig | 353090 | 2886 | 5.53E-03 | 3.65E-02 | | 5.72E-03 | | 3.24E-04 | | | | | | | |
| *Celetes* | full | 359439 | 0 | 1.57E-02 | 3.06E-03 | | 2.67E-03 | | 4.34E-03 | | 1.78 | 0.88 | | | | |
| *decolor* OTU 1 | no mig | 363095 | 3656 | 2.94E-02 | 3.99E-03 | | 5.88E-03 | | 1.26E-03 | | | | | | | |
| *Celetes* | full | 99728 | 0 | 3.52E-03 | 4.66E-03 | | 6.47E-04 | | 9.29E-03 | | 3.37 | 3.73 | | | | |
| *decolor* OTU 2 | no mig | 99910 | 182 | 3.69E-03 | 6.95E-03 | | 5.99E-03 | | 3.19E-04 | | | | | | | |
| *Celetes impar* | full | 644259 | 0 | 7.44E-03 | 3.49E-03 | 6.66E-03 | 1.98E-03 | 3.50E-04 | 4.33E-03 | 8.89E-03 | 0.08 | 0.13 | 0.61 | 2.14 | 0.38 | 0.23 |
| | noAC | 645339 | 1080 | 7.43E-03 | 3.46E-03 | 6.58E-03 | 1.98E-03 | 4.53E-04 | 4.33E-03 | 9.19E-03 | | | 1.04 | 1.97 | 0.57 | 0.28 |
| | no A,B | 646163 | 1904 | 7.89E-03 | 3.51E-03 | 6.08E-03 | 1.07E-03 | 1.14E-03 | 6.53E-03 | 1.95E-03 | 0.08 | 0.3 | 0.53 | 2.69 | | |
| | no A | 649068 | 4809 | 7.11E-03 | 3.36E-03 | 6.12E-03 | 1.33E-03 | 2.83E-03 | 4.79E-03 | 1.40E-03 | | | 1.18 | 2.24 | | |
| | no B,C | 650212 | 5952 | 7.14E-03 | 3.48E-03 | 8.77E-03 | 1.16E-02 | 2.06E-03 | 8.81E-04 | 5.42E-03 | 0.14 | 0.12 | | | 0.68 | 0.41 |
| | no C | 651627 | 7368 | 7.14E-03 | 3.46E-03 | 7.21E-03 | 1.27E-02 | 2.93E-03 | 7.98E-04 | 4.77E-03 | | | | | 0.9 | 0.25 |
| | no B | 654584 | 10325 | 5.94E-03 | 3.56E-03 | 9.38E-03 | 1.25E-02 | 3.67E-03 | 7.77E-04 | 4.17E-03 | 0.04 | 0.8 | | | | |
| | no mig | 657909 | 13649 | 6.96E-03 | 4.01E-03 | 9.67E-03 | 9.76E-03 | 6.06E-03 | 9.19E-04 | 2.17E-03 | | | | | | |
| *Dialomia polyphaga* | full | 107341 | 0 | 4.30E-03 | 3.39E-03 | | 8.61E-04 | | 5.18E-03 | | 0.34 | 0.59 | | | | |
| | no mig | 108242 | 902 | 5.89E-03 | 3.06E-03 | | 1.83E-03 | | 1.32E-03 | | | | | | | |
| *Microstrates* | full | 135833 | 0 | 1.47E-02 | 3.46E-03 | | 3.60E-04 | | 8.04E-03 | | 0.17 | 0.24 | | | | |
| *bondari* OTU 1 | no mig | 136929 | 1096 | 1.68E-02 | 3.76E-03 | | 1.09E-03 | | 3.63E-03 | | | | | | | |
| *Microstrates ypsilon* | full | 218442 | 0 | 1.06E-02 | 4.48E-03 | | 1.67E-03 | | 5.64E-03 | | 1.27 | 1.06 | | | | |
| | no mig | 221062 | 2620 | 1.34E-02 | 5.77E-03 | | 5.98E-03 | | 1.25E-03 | | | | | | | |
| *Remertus rectinasus* | no B,C | 112239 | 0 | 3.64E-03 | 3.24E-03 | 3.75E-03 | 4.72E-04 | 1.11E-04 | 2.96E-03 | 1.14E-02 | 0.31 | 0.03 | 3.82 | 1.57 | | |
| | full | 112254 | 15 | 6.12E-03 | 3.46E-03 | 4.42E-03 | 5.16E-04 | 8.86E-05 | 2.65E-03 | 1.10E-02 | 0.49 | 0 | 8.03 | 0.39 | 0.02 | 0.07 |
| | no A,B | 112307 | 68 | 7.66E-03 | 3.50E-03 | 3.70E-03 | 6.36E-04 | 6.90E-05 | 1.09E-02 | 1.90E-03 | | | 12.01 | 0 | 0.08 | 0.04 |
| | noAC | 112372 | 134 | 1.03E-02 | 3.24E-03 | 5.36E-03 | 1.20E-03 | 3.93E-04 | 1.04E-03 | 1.04E-02 | 0.61 | 0 | | | 0.06 | 0.11 |
| | no B | 112878 | 639 | 7.25E-03 | 3.51E-03 | 3.18E-03 | 9.98E-04 | 3.73E-04 | 1.07E-02 | 1.07E-03 | | | 18.81 | 0.47 | | |
| | no A | 113038 | 800 | 2.57E-02 | 3.46E-03 | 4.37E-03 | 4.62E-04 | 8.64E-05 | 1.31E-03 | 1.08E-02 | | | | | 0.08 | 0.07 |
| | no C | 113224 | 986 | 4.76E-03 | 3.24E-03 | 6.60E-03 | 4.11E-03 | 7.35E-03 | 5.86E-04 | 5.97E-03 | 0.41 | 0.1 | | | | |
| | no mig | 113631 | 1392 | 6.84E-03 | 3.06E-03 | 5.05E-03 | 1.79E-03 | 5.05E-03 | 1.04E-03 | 4.26E-03 | | | | | | |

Table S2: Model choice by cross validation in BEDASSLE2. For each OTU, models are ordered by average predictive value. The preferred model (in bold) is the one including fewer parameters among those with highest posterior predictive value, considering the 95% confidence intervals. Models are labelled after the distance matrices included: geographical distance (*g*), plant host genetic distance (*p*) and climatic distance (*c*). In the null model, genetic covariance across samples does not depend deterministically on any other distance matrix.

| OTU | Model | Predictive value, mean (95% confidence interval) |
|---|---|---|
| *Anchylorhynchus trapezicollis* OTU 1 | **gp** | **81987.80(81987.69–81987.92)** |
| | gpc | 81987.52(81987.42–81987.63) |
| | pc | 81982.12(81982.02–81982.22) |
| | p | 81982.07(81981.97–81982.17) |
| | gc | 80228.69(80228.58–80228.80) |
| | c | 78185.06(78184.95–78185.18) |
| | g | 77827.47(77827.39–77827.55) |
| | null | 69095.10(69095.01–69095.20) |
| *Anchylorhynchus trapezicollis* OTU 2 | gp | 50557.87(50557.77–50557.97) |
| | gpc | 50557.84(50557.77–50557.92) |
| | **g** | **50557.78(50557.71–50557.86)** |
| | gc | 50557.75(50557.67–50557.83) |
| | pc | 49903.85(49903.76–49903.94) |
| | p | 49903.74(49903.65–49903.83) |
| | c | 49058.37(49058.28–49058.46) |
| | null | 46977.82(46977.73–46977.91) |
| *Anchylorhynchus trapezicollis* OTU 3 | **gp** | **1467.62(1467.60–1467.64)** |
| | gpc | 1467.58(1467.56–1467.60) |
| | p | 1467.40(1467.38–1467.42) |
| | g | 1467.33(1467.31–1467.35) |
| | pc | 1467.26(1467.24–1467.28) |
| | gc | 1467.15(1467.14–1467.17) |
| | c | 1466.17(1466.15–1466.18) |
| | null | 1458.93(1458.92–1458.95) |
| *Andranthobius bondari* OTU 1 | **g** | **70595.05(70594.95–70595.14)** |
| | gp | 70595.02(70594.93–70595.12) |
| | gc | 70595.00(70594.91–70595.09) |
| | gpc | 70594.67(70594.58–70594.75) |
| | p | 70581.85(70581.76–70581.95) |
| | pc | 70581.83(70581.75–70581.90) |
| | c | 70579.11(70579.02–70579.21) |
| | null | 70571.69(70571.59–70571.78) |
| *Andranthobius bondari* OTU 2 | **p** | **29694.04(29693.95–29694.12)** |
| | pc | 29693.86(29693.78–29693.94) |
| | gp | 29693.75(29693.68–29693.82) |
| | gpc | 29693.72(29693.65–29693.79) |
| | g | 29690.56(29690.49–29690.63) |
| | c | 29690.13(29690.05–29690.20) |
| | gc | 29688.93(29688.87–29688.99) |
| | null | 29646.84(29646.76–29646.93) |
| *Celetes decolor* OTU 1 | **g** | **19118.70(19118.64–19118.76)** |
| | gc | 19118.70(19118.65–19118.74) |
| | gp | 19118.67(19118.62–19118.72) |
| | gpc | 19118.49(19118.44–19118.54) |
| | p | 18925.93(18925.87–18925.98) |
| | pc | 18916.29(18914.95–18917.63) |
| | c | 18751.01(18750.95–18751.06) |
| | null | 18030.67(18030.61–18030.73) |

Table S2: (continued)

| Species | Model | Predictive value, mean (95% confidence interval) |
|---|---|---|
| *Celetes decolor* OTU 2 | **g** | **3423.65(3423.63–3423.68)** |
| | gp | 3423.55(3423.53–3423.58) |
| | gpc | 3423.51(3423.48–3423.55) |
| | gc | 3423.28(3423.21–3423.35) |
| | c | 3409.73(3409.59–3409.88) |
| | pc | 3408.21(3407.67–3408.75) |
| | p | 3392.80(3392.76–3392.84) |
| | null | 3299.74(3299.71–3299.76) |
| *Celetes impar* | gpc | 87977.06(87976.94–87977.18) |
| | **gp** | **87977.02(87976.92–87977.13)** |
| | gc | 87973.52(87973.40–87973.64) |
| | g | 87973.40(87973.28–87973.51) |
| | pc | 87621.84(87621.74–87621.95) |
| | p | 87621.82(87621.72–87621.91) |
| | c | 86979.59(86979.48–86979.70) |
| | null | 83617.98(83617.87–83618.09) |
| *Dialomia polyphaga* | gc | 7679.66(7679.63–7679.70) |
| | **g** | **7679.64(7679.60–7679.67)** |
| | gp | 7679.61(7679.57–7679.64) |
| | gpc | 7679.50(7679.47–7679.53) |
| | p | 7632.07(7632.00–7632.14) |
| | pc | 7631.79(7631.72–7631.86) |
| | c | 7621.51(7621.35–7621.66) |
| | null | 7579.66(7579.63–7579.70) |
| *Microstrates bondari* OTU 1 | **gp** | **13190.15(13190.07–13190.22)** |
| | gpc | 13189.95(13189.89–13190.01) |
| | g | 13182.32(13182.28–13182.36) |
| | gc | 13182.28(13182.24–13182.33) |
| | pc | 13177.53(13177.49–13177.58) |
| | p | 13172.54(13172.50–13172.59) |
| | c | 13145.76(13145.61–13145.92) |
| | null | 13102.05(13102.01–13102.10) |
| *Microstrates ypsilon* | **gp** | **18338.82(18338.77–18338.87)** |
| | gpc | 18338.76(18338.72–18338.81) |
| | g | 18337.02(18336.97–18337.07) |
| | gc | 18336.90(18336.85–18336.96) |
| | p | 18253.96(18253.92–18254.00) |
| | pc | 18253.94(18253.88–18253.99) |
| | c | 18051.73(18051.67–18051.78) |
| | null | 17820.16(17820.10–17820.21) |
| *Phytotribus cocoseae* | **gp** | **22587.74(22587.67–22587.81)** |
| | gpc | 22587.68(22587.61–22587.75) |
| | g | 22584.24(22584.17–22584.30) |
| | gc | 22584.21(22584.15–22584.27) |
| | pc | 22569.90(22569.84–22569.96) |
| | c | 22563.24(22563.18–22563.30) |
| | p | 22537.11(22537.00–22537.21) |
| | null | 22417.75(22417.69–22417.80) |
| *Remertus rectinasus* | **gp** | **10848.63(10848.59–10848.67)** |
| | gpc | 10848.60(10848.56–10848.64) |
| | p | 10843.44(10843.40–10843.48) |
| | pc | 10843.41(10843.36–10843.45) |
| | gc | 10393.50(10393.46–10393.53) |
| | c | 10167.43(10167.40–10167.47) |
| | g | 9933.43(9933.39–9933.47) |
| | null | 7643.13(7640.30–7645.96) |

Table S3: Mean and 95% credibility intervals for parameters estimated by the best BEDASSLE 2 model for each weevil OTU. $\alpha_2$ controls the shape of genetic covariance decay, $\gamma$ is the average covariance across samples and $\alpha_0$ is the covariance at distance 0. The other $\alpha$ parameters control the contribution of geography or plant host.

| plant host | Interaction | species | $\alpha_{geo}$ | $\alpha_{plant}$ | $\alpha_0$ | $\alpha_2$ |
|---|---|---|---|---|---|---|
| *S. botryophora* | pollinator,antagonist | *Anc. trapezicollis* OTU 3 | 1.62e (0.43–2.49)e-01 | 1.58 (0.45–2.43)e-01 | 0.06 (0.06–0.07) | 1.04 (0.36–1.87) |
| | commensal | *Andranthobius bondari* OTU 2 | 0.78 (0.02–2.94)e-03 | 1.14 (0.08–3.33)e-02 | 0.18 (0.18–0.19) | 1.20 (0.72–1.80) |
| | commensal | *Celetes decolor* OTU 2 | 2.14 (2.01–2.26)e-01 | 1.28 (0.06–3.49)e-02 | 0.14 (0.13–0.15) | 1.93 (1.76–2.00) |
| | commensal | *Phytotribus cocoseae* | 4.40 (3.19–5.58)e-02 | 2.64 (1.49–4.01)e-02 | 0.18 (0.18–0.19) | 1.67 (1.41–1.95) |
| | antagonist | *Microstrates bondari* OTU 1 | 5.52 (4.41–6.20)e-02 | 3.85 (2.88–4.40)e-02 | 0.20 (0.19–0.21) | 1.90 (1.69–2.00) |
| *S. coronata* | pollinator,antagonist | *Anc. trapezicollis* OTU 2 | 5.25 (4.93–5.54)e-02 | 1.95 (0.06–5.36)e-04 | 0.20 (0.19–0.20) | 1.03 (1.01–1.05) |
| | commensal | *Andranthobius bondari* OTU 1 | 0.92 (0.22–2.37)e-04 | 1.50 (0.03–6.54)e-06 | 0.22 (0.22–0.23) | 0.74 (0.64–0.85) |
| | commensal | *Celetes decolor* OTU 1 | 7.01 (6.46–7.57)e-02 | 0.36 (0.01–1.04)e-03 | 0.20 (0.19–0.20) | 1.13 (1.09–1.17) |
| | commensal | *Celetes impar* | 6.64 (5.90–7.47)e-03 | 2.54 (1.91–3.18)e-04 | 0.22 (0.21–0.23) | 0.66 (0.64–0.67) |
| | commensal | *Dialomia polyphaga* | 0.92 (0.48–1.54)e-02 | 0.42 (0.01–1.42)e-04 | 0.20 (0.19–0.21) | 0.43 (0.38–0.50) |
| | antagonist | *Microstrates ypsilon* | 3.92 (3.52–4.32)e-02 | 5.57 (4.12–7.09)e-03 | 0.21 (0.20–0.22) | 1.32 (1.27–1.37) |
| both | pollinator,antagonist | *Anc. trapezicollis* OTU 1 | 1.96 (1.74–2.17)e-02 | 1.61 (1.57–1.66)e-01 | 0.23 (0.22–0.23) | 1.76 (1.73–1.78) |
| | antagonist | *Remertus rectinasus* | 6.40 (5.70–7.06)e-02 | 3.59 (3.50–3.67)e-01) | 0.20 (0.19–0.21) | 2.00 (2.00–2.00) |

Table S4: Number of samples, populations and SNPs sequenced for each plant species and weevil OTU, following filtering for HWE equilibrium. For weevil OTUs, we also include the number of unlinked SNPs used in BEDASSLE, filtered as described in the main text methods.

| Species/OTU | Samples | Populations | All SNPs | variable RAD loci | Unlinked SNPs |
|---|---|---|---|---|---|
| *Syagrus botryophora* | 25 | 5 | 12532 | 5300 | – |
| *Syagrus coronata* | 49 | 13 | 30089 | 10081 | – |
| *Anchylorhynchus trapezicollis* OTU 1 | 47 | 13 | 49408 | 12829 | 8607 |
| *Anchylorhynchus trapezicollis* OTU 2 | 36 | 10 | 38022 | 12558 | 8520 |
| *Anchylorhynchus trapezicollis* OTU 3 | 5 | 3 | 7799 | 4097 | 3008 |
| *Andranthobius bondari* OTU 1 | 38 | 12 | 51948 | 11724 | 9098 |
| *Andranthobius bondari* OTU 2 | 22 | 5 | 32082 | 11614 | 8798 |
| *Celetes decolor* OTU 1 | 20 | 8 | 31706 | 9505 | 6206 |
| *Celetes decolor* OTU 2 | 9 | 4 | 9596 | 4321 | 3286 |
| *Celetes impar* | 47 | 13 | 56309 | 11628 | 9786 |
| *Dialomia polyphaga* | 18 | 9 | 10542 | 4511 | 2794 |
| *Microstrates bondari* OTU 1 | 23 | 5 | 15973 | 3984 | 3475 |
| *Microstrates ypsilon* | 27 | 11 | 21256 | 5345 | 3838 |
| *Phytotribus cocoseae* | 26 | 5 | 15305 | 6017 | 5695 |
| *Remertus rectinasus* OTU 1 | 18 | 8 | 14371 | 5006 | 3694 |

#### References


[1] GBIF.org. *GBIF Ocurrence Download.* 2019. DOI: 10.15468/dl.lprfwo. URL: https://doi.org/10.15468/dl.lprfwo.

[2] Larry R. Noblick. “A revision of the genus Syagrus (Arecaceae).” In: *Phytotaxa* 294.1 (2017), pp. 001–262. DOI: 10.11646/phytotaxa.294.1.1.
